# AlphaFold Prediction of Structural Ensembles of Disordered Proteins

**DOI:** 10.1101/2023.01.19.524720

**Authors:** Z. Faidon Brotzakis, Shengyu Zhang, Michele Vendruscolo

## Abstract

Deep learning methods of predicting protein structures have reached an accuracy comparable to that of high-resolution experimental methods. It is thus possible to generate accurate models of the native states of hundreds of millions of proteins. An open question, however, concerns whether these advances can be translated to disordered proteins, which should be represented as structural ensembles because of their heterogeneous and dynamical nature. Here we show that the inter-residue distances predicted by AlphaFold for disordered proteins can be used to construct accurate structural ensembles. These results illustrate the application to disordered proteins of deep learning methods originally trained for predicting the structures of folded proteins.

## Introduction

The application of deep learning methods to the protein folding problem has transformed our ability to generate accurate models of the native states of proteins from the knowledge of their amino acid sequences^1-3^. Furthermore, the initial predictions of the native structures of proteins have been recently extended to protein complexes^4,5^.

These advances have prompted the question of whether it is possible to use this type of approach for the prediction of the conformational fluctuations of the native states of folded proteins^6-12^, and more generally for the characterisation of the structural properties of the native states of disordered proteins^13-15^. Support for this idea comes from the observation that AlphaFold performs as well as current state-of-the-art predictors of protein disorder^16,17^. It has also been reported that the predicted aligned error (PAE) maps from AlphaFold are correlated with the distance variation matrices from molecular dynamics simulations^7^, suggesting that AlphaFold provides information about the dynamics of proteins in addition to their structures.

Since the native states of disordered proteins can be represented in terms of ensembles of conformations with statistical weights obeying the Boltzmann distribution^13,18^, the task is to extend AlphaFold to predict structural ensembles. In this work, we propose an approach to perform this task. We based this approach on the observation that AlphaFold can predict inter-residue distances even for disordered proteins, despite having been trained on folded proteins. This is important because it enables the transfer of inter-residue distance information from folded proteins, whose accurate structures are available in large numbers in the Protein Data Bank (PDB)^19^ for the training of deep learning methods, to disordered proteins, whose structural ensembles have been determined in much lower numbers and with less accuracy^20^.

The inter-residue distances predicted by AlphaFold are collected together in the form of a distance map, from which the structure of a protein can be constructed^1^. For a disordered protein, the construction problem becomes to translate the predicted distance map into a structural ensemble, rather than a single structure. There are many ways to achieve this objective. For example, the predicted distances can be used as structural restraints in molecular dynamics simulations^18,21,22^. Here, we adopt a strategy based on the reweighting of a pre-calculated structural ensemble^23,24^. In this way, the weights of the different structures in the ensemble are fine tuned to make the reweighted ensemble maximally consistent with the inter-residue distances provided by AlphaFold. We illustrate the approach using the examples of structural ensembles of Aβ^25^, a disordered peptide closely associated with Alzheimer’s disease^26,27^, and of α-synuclein^28^, a disordered protein whose aggregation in Lewy bodies is the hallmark of Parkinson’s disease^29^.

## Results

### Structural ensemble of Aβ

To use AlphaFold to determine a structural ensemble representing the native state of Aβ, we started from a recently reported structural ensemble^25^ (i.e. the prior ensemble, referred here as the MD ensemble of Aβ) which we adopted as starting point for a reweighting procedure^23^ (see Methods). A comparison of the inter-residue distances back-calculated from the MD ensemble and those predicted by AlphaFold is shown in **Figure 1A**. Since the MD ensemble of Aβ is in good agreement with experimental data^25^, the correlation reported in **Figure 1A** indicates that AlphaFold can predict inter-residue distances in the native state of Aβ, which is a disordered peptide.

**Figure 1.**
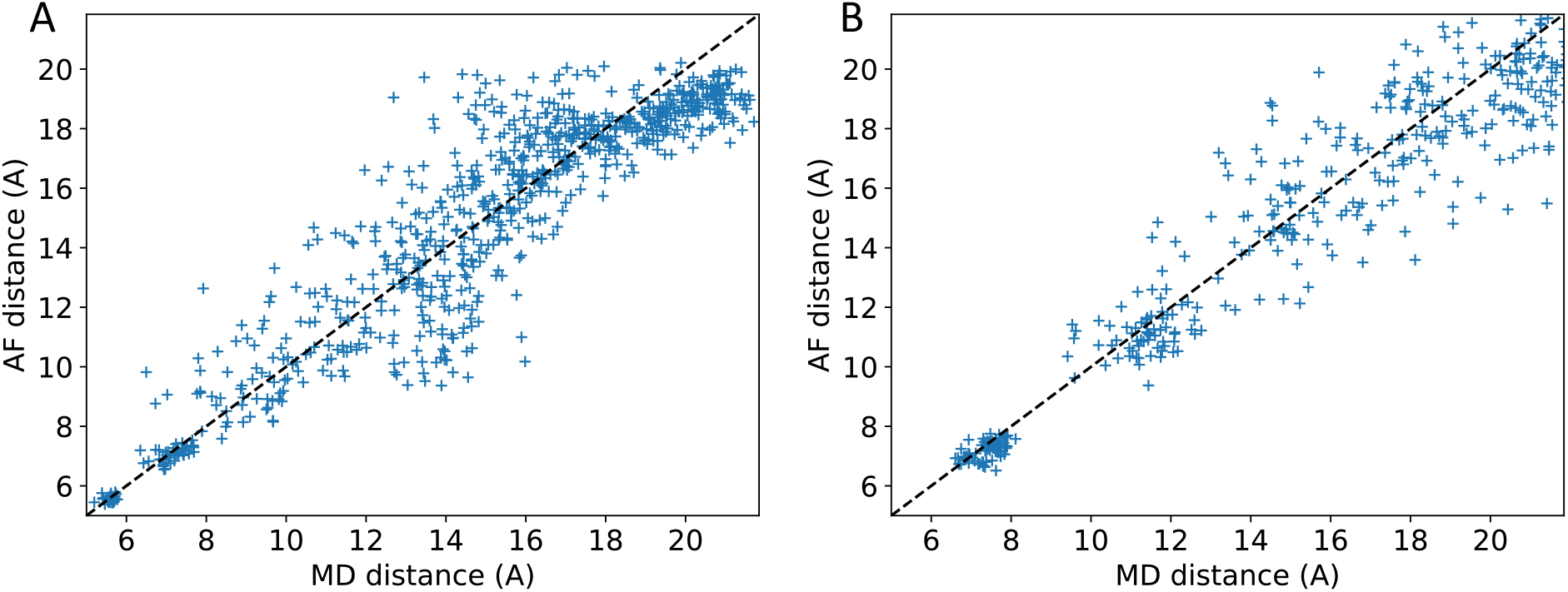
Comparison of the inter-residue distances back-calculated from the MD ensembles and those predicted by AlphaFold. **(A**,**B)** Inter-residue distances back-calculated from the Aβ **(A)** and α-synuclein **(B)** MD ensembles are reported on the x-axis, and those predicted by AlphaFold on the y-axis. The cut-off distance in the AlphaFold predictions is 21.84 Å.

The application of the reweighting procedure (see Methods) resulted in the generation of a posterior ensemble, which we call the AF-MD ensemble of Aβ (**Figure 2A,B**). The reweighting procedure changed the statistical weights of the structures in the MD ensemble to achieve a closer match between the inter-residue distances predicted by AlphaFold and those back-calculated from the AF-MD ensemble (**Figure S1A**). The overall *χ*^2^ was reduced from 0.26 in the MD ensemble to 0.02 in the AF-MD ensemble. The root mean square distance (RMSD) between distances was reduced from 1.97 Å in the MD to 0.52 Å in the AF-MD ensemble. The number of distance violations, which are defined as distances in the AF ensemble that are not within the AF error from the AF distance average, is reduced from 3 in the MD ensemble to 0 in the AF-MD ensemble.

**Figure 2.**
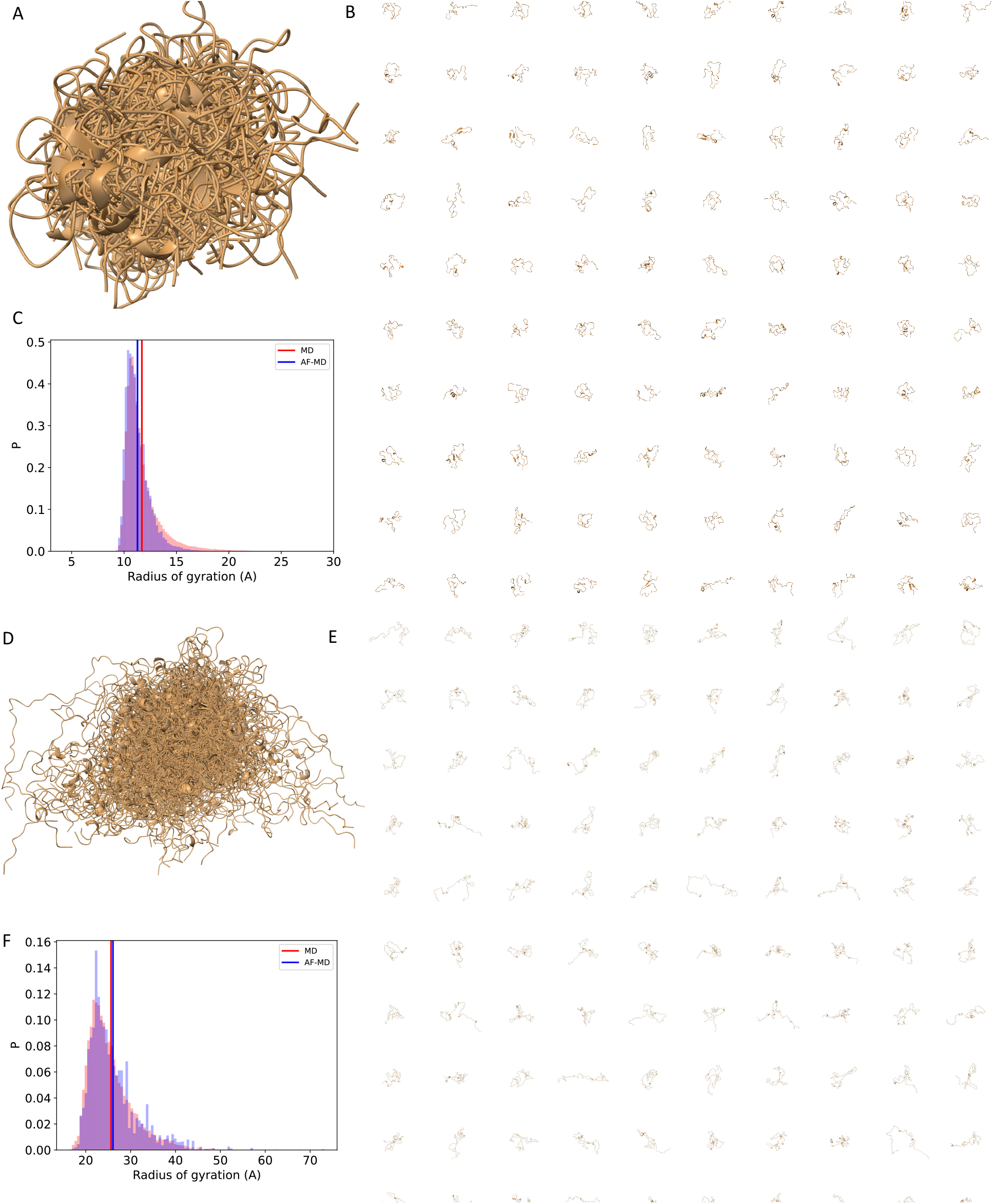
Structural ensembles of Aβ and α-synuclein predicted by AlphaFold using molecular dynamics to generate the prior ensembles. **(A**,**D)** Illustration of the structural ensemble. (**B**,**E**) Individual 100 representative structures. (**C**,**F**) Distributions of the tadius of gyration in the MD ensemble (pink) and AF-MD ensemble (purple); the average values are shown in red and blue solid lines, respectively.

A comparison of the structural properties of the MD and AF-MD ensembles is shown in **Figures 2C and S2**,**3**. Both the secondary structure populations (**Figure S3A-C**) and the distribution of the radius of gyration (**Figure 1C**) did not change significantly, as expected given the correlation between inter-residue distances reported in **Figure 1A**. The secondary structure of Aβ in the AF-MD ensemble was slightly more β-sheet like, less α-helical (**Figure S3A-C**) compared to the MD ensemble. The average radius of gyration was 11.6 ± 1.9 Å and 11.2 ± 1.3 Å in the MD and AF-MD ensembles, respectively.

Although accurate, molecular dynamics are slow in generating prior ensembles due to very large size of the conformational space accessible to disordered proteins. To overcome this problem, we explored the use faster approaches based on deep learning. We present the results obtained with FoldingDiff, a denoising diffusion probabilistic model trained with folded protein structures^30^. We first generated a prior structural ensemble of backbones of protein length of 42 (see Methods), which we refer to as FD ensemble of Aβ. We then utilized it as a starting point for the reweighting procedure described above. A comparison of the inter-residue distances back-calculated from the FD ensemble and those predicted by AlphaFold is shown in **Figure 3A**. The FD ensemble is a prior of lower quality than the MD ensemble, showing *x*^2^ Of the constrained distances of 1.34 compared to 0.26 of the MD ensemble. However, one can still use the FD ensemble in a reweighting procedure to produce the AF-FD ensemble of Aβ (**Figure 3B,C**).

**Figure 3.**
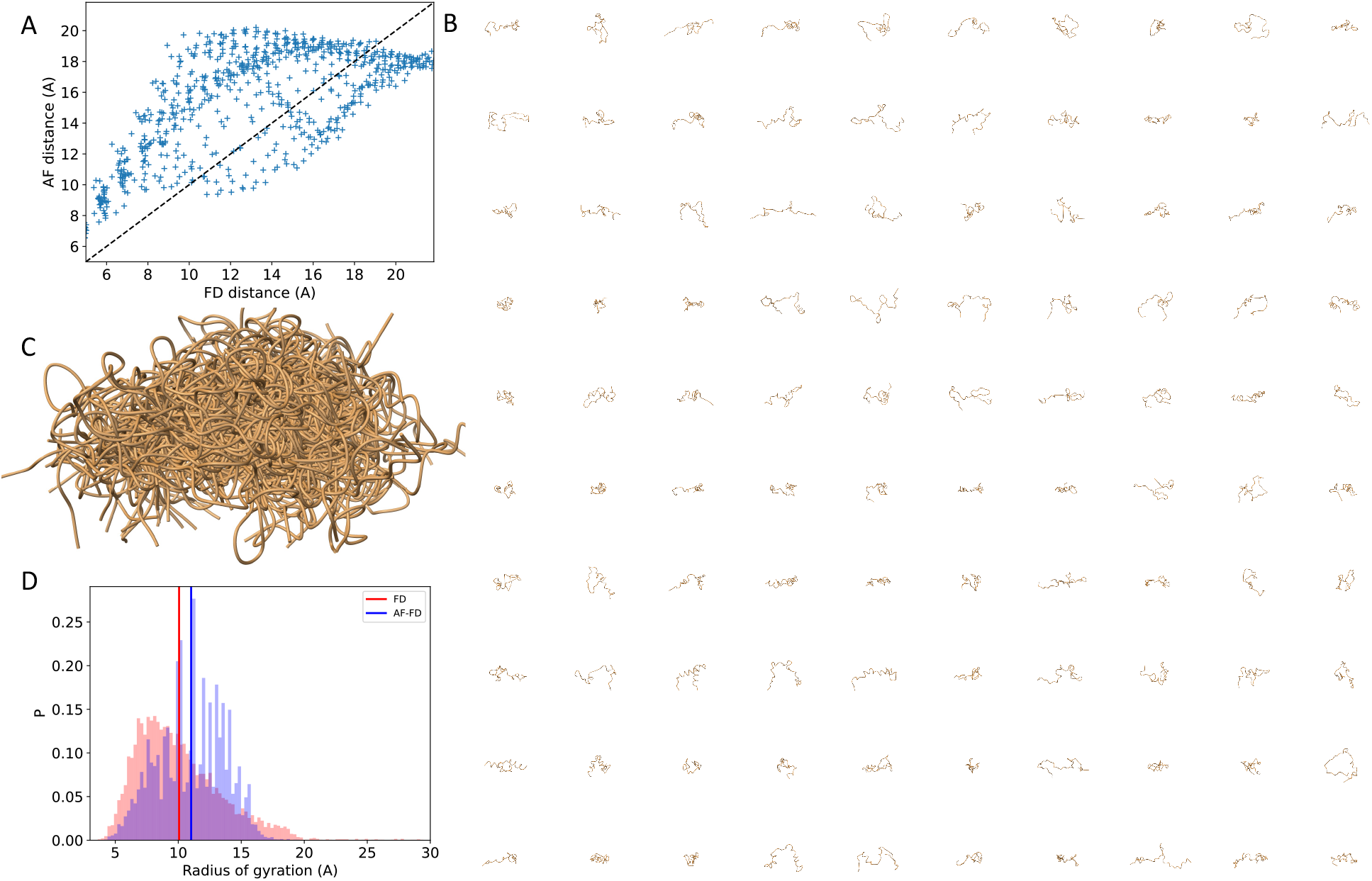
Structural ensemble of Aβ predicted by AlphaFold using FoldingDiff to generate the prior ensemble. **(A)** Comparison of the inter-residue distances calculated from the FD ensemble and those predicted by AlphaFold for Aβ. **(B**,**C)** 100 representative structures of the AF-FD ensemble. **(D)** Distribution of the radius of gyration in the FD ensemble (pink) and AF-FD ensemble (purple); the average values are shown in red and blue solid lines, respectively.

The reweighting procedure changed the statistical weights of the structures in the FD ensemble to achieve a closer match between the inter-residue distances predicted by AlphaFold and those back-calculated from the FD ensemble (**Figure S1C**). The overall *χ*^2^ was reduced to 0.23 in the AF-FD ensemble. The RMSD was reduced from 4.07 Å in the FD ensemble to 1.71 Å in the AF-FD ensemble. The number of distance violations was reduced from 25 in the FD ensemble to 1 in the AF-FD ensemble.

A comparison of the structural properties of the FD and AF-FD ensembles is shown in **Figure 3D**. The average radius of gyration value changes from 10.0 ± 3.4 Å to 11.0 ± 2.7 Å from the FD to the AF-FD ensembles. The average value of the radius of gyration of the AF-FD ensemble is close to that of the AF-MD ensemble despite the fact that the FD ensemble contains more compact structures than the MD ensemble (**Figures 2C and 3D**). This is consistent with previous finding that reweighting with many constraints produces posterior ensembles of high quality even from prior ensembles of relatively quality^31^.

### Structural ensemble of α-synuclein

To use AlphaFold to determine a structural ensemble representing the native state of α-synuclein, we started from a recently reported structural ensemble^28^ as prior ensemble, referred here as the MD ensemble of α-synuclein. We used this ensemble as starting point for the same reweighting procedure described above for the MD ensemble of Aβ. The comparison of the inter-residue distances back-calculated from the MD ensemble and those predicted by AlphaFold is shown in **Figure 1B**. Since the MD ensemble of α-synuclein is in good agreement with experimental data^28^, the correlation reported in **Figure 1B** indicates that AlphaFold can predict inter-residue distances even in the native state of α-synuclein, which is a disordered protein.

The application of the reweighting procedure generated a posterior structural ensemble, called the AF-MD ensemble of α-synuclein (**Figure 2D,E**). As in the case of Aβ, the reweighting procedure changed the statistical weights of the structures in the MD ensemble to achieve a closer match between the inter-residue distances predicted by AlphaFold and those back-calculated from the AF-MD ensemble (**Figure S1B**). The overall *χ*^2^ is reduced from 0.26 in the MD ensemble to 0.02 in the AF-MD ensemble. The overall *RMSD*reduces from 1.97 Å in the prior-MD to 0.52 Å in the AF ensemble. The number of distance violations is reduced from 3 in the MD ensemble to 0 in the AF-MD ensemble.

A comparison of the structural properties of the MD and AF-MD ensembles of α-synuclein is shown in **Figures 2F, S2C-D and S3D-F**. Both the secondary structure populations (**Figure S3D-F**) and the distribution of the radius of gyration (**Figure 2F**) did not change significantly, as expected given the correlation between inter-residue distances reported in **Figure 1B**. The secondary structure of α-synuclein in the AF-MD ensemble was slightly more α-helical and less β-sheet like (**Figure S3D-F**) compared to the MD ensemble. The average radius of gyration was 25.6 ± 5.7 Å and 26.2 ± 5.9 Å in the MD and AF-MD ensembles, respectively.

## Discussion and Conclusions

We have described a method of using AlphaFold to generate structural ensembles representing the native states of disordered proteins. The method is based on the observation that the inter-residue distances predicted by AlphaFold can be used as structural restraints to construct structural ensembles (**Figures 1**).

The finding that AlphaFold may be able to predict inter-residue contacts even for disordered proteins, despite the fact that disordered proteins are not present in the AlphaFold training dataset, is perhaps surprising. Since deep learning methods do not readily generalise to cases not observed during training, this finding could indicate that the types of interactions between residues that stabilise the native states of disordered proteins do not differ fundamentally from those that stabilise the native states of folded proteins, but possibly they are just weaker. In any case, we anticipate that the systematic inclusion of experimentally-derived structural ensembles of proteins^20^, as they will become available in the future, in the training of deep learning methods for the prediction of protein structures will improve the accuracy of the predictions of structural ensembles of disordered proteins.

To generate structural ensembles, in the reweighting approach adopted here, we have used as starting point structural ensembles obtained by all-atom molecule dynamics simulations. This approach is rather accurate, but time consuming^25,28^. Alternative methods based on the translation of the distance map predicted by AlphaFold into structural ensembles using the distances as structural restraints in all atom molecular dynamics simulations^18,21,22^ could be equally time consuming. Faster results can be obtained by generating initial ensembles using methods based on coarse graining^32,33^ or deep learning^30,34^, and then relying on reweighting procedures to obtain more accurate statistical weights.

Overall, the results that we have presented illustrate the use of deep learning methods originally developed for predicting the native states of folded proteins^1-3^ to generate structural ensembles representing the native states of disordered proteins, or of proteins containing disordered regions. The scope of protein structure predictions based on deep learning could thus be considerably extended.

## Methods

### AlphaFold

Inter-residue distances of disordered proteins were predicted through the distogram head of AlphaFold^1^. These distances are defined as those between the β carbon atom positions for all amino acids except glycine, for which the α carbon atom positions were instead used. The multiple sequence alignment (MSA) was conducted by MMseqs2^35^ (default setting) on BFD/MGnify^3^ and Uniclust30^36^. No structural template was used for the prediction. Model 1.1.1 of AlphaFold (default setting)^1^ was used for the predictions, and no structural template was used.

AlphaFold describes the distribution of inter-residue distances into 64 bins of equal width, covering the range from 2 to 22 Å, with the last bin also including distances longer than 22 Å. For each pair of residues (*i* and *j*), AlphaFold predicts the probability 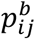 that their distance is within bin *b*. The predicted distance 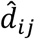 and the standard deviation *σ*_*ij*_ of the predicted distribution of the distances between residue *i* and *j* are calculated by

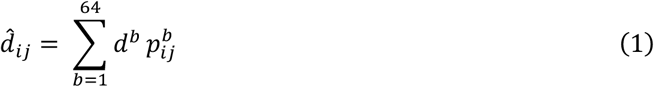

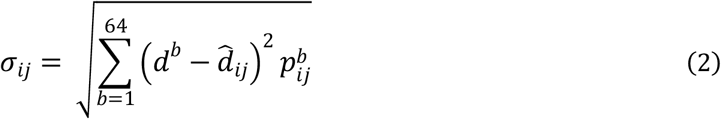

where *d*^*d*^ represents the central value of bin *b*, which ranges from 2.15625 to 21.84375 Å.

### Reweighting

A reweighting procedure starts from a pre-calculated structural ensemble of *n* structures ***X*** = {*X*_1_, *X*_2_,.., *X*_*n*_}, which is referred to as prior ensemble. The prior ensemble is characterised by a prior distribution *P*^0^(***X***) where each structure has an associated statistical weight 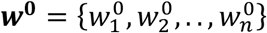. *P*^0^(***X***) can be generated with physics based methods, such as molecular dynamics^25,28^, or with machine learning methods such as generative models^30,34^. However, various approximations in the generation procedure, for example due to inaccurate force field in molecular dynamics or to low-quality structural data in the training of neural networks, imply that *P*^0^(***X***) might differ from the true distribution of structures *P*(***X***). The distance between *P*^0^(***X***) and *P*(***X***) can be quantified by the Shannon entropy

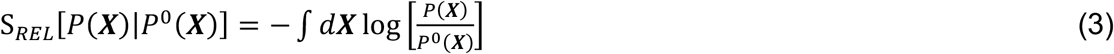

where the right-hand side is the negative of the Kullback-Leibler divergence between distributions.

The central aim of the reweighting procedure is to modify the statistical weights of the prior distribution in order to obtain a new distribution, *P*^1^(***X***), known as the posterior distribution, which minimises S_*REL*_ for a given *P*(***X***). The reweighting is carried out by using information about *P*(***X***).

In the context of this study, for a protein of *N* amino acids, the information about *P*(***X***) is provided by an *N x N* distance matrix ***d***^*AF*^ between amino acids, which is obtained using AlphaFold. From the prior distribution *P*^0^(***X***), the distances between amino acids can be calculated as weighted averages, giving a prior distance matrix ***d***^0^. ***d***^*AF*^ is used as constraints to reweight *P*^0^(***X***) to generate *P*^1^(***X***), which corresponds to a posterior distance matrix ***d***^1^, so that the difference, denoted as *χ*^2^, between ***d***^1^ and ***d***^*AF*^ is minimised. To achieve this goal we use the Bayesian/Maximum Entropy (BME) approach^37^ with θ=0.5 and 1 for Aβ and α-synuclein, respectively, for reweighting the MD ensembles, and θ=4 for reweighting the Aβ FD ensemble. These values generated posterior ensembles substantially increasing the overall *χ*^2^ agreement with the AlphaFold distance constraints. In practice, rather than constraining all distances, we did not include neighbouring residues (up to 4 residues) for Aβ and that have a distance larger than 21.8 Å in the MD or FD ensembles. For the α-synuclein calculations we do not include neighbouring residues (up to 2 residues) and that have a distance larger than 21.8 Å in the MD ensemble.

### FoldingDiff

To calculate the prior ensemble of Aβ using FoldingDiff, we used the pre-trained FoldingDiff model to generate a prior structural ensemble of backbone conformations for a polypeptide chain of 42 residues, following the approach in the FoldingDiff github repository^30^. We generated 100 structures while keeping the full folding history, thus resulting in an ensemble of 75000 structures. This calculation was rather fast, lasting 2 hours using 6 CPUs. We note that we did not repeat the calculations to generate a prior ensemble for α-synuclein since the current implementation of FoldingDiff has a maximum length of 127 amino acids.

---

**Figure S1.**
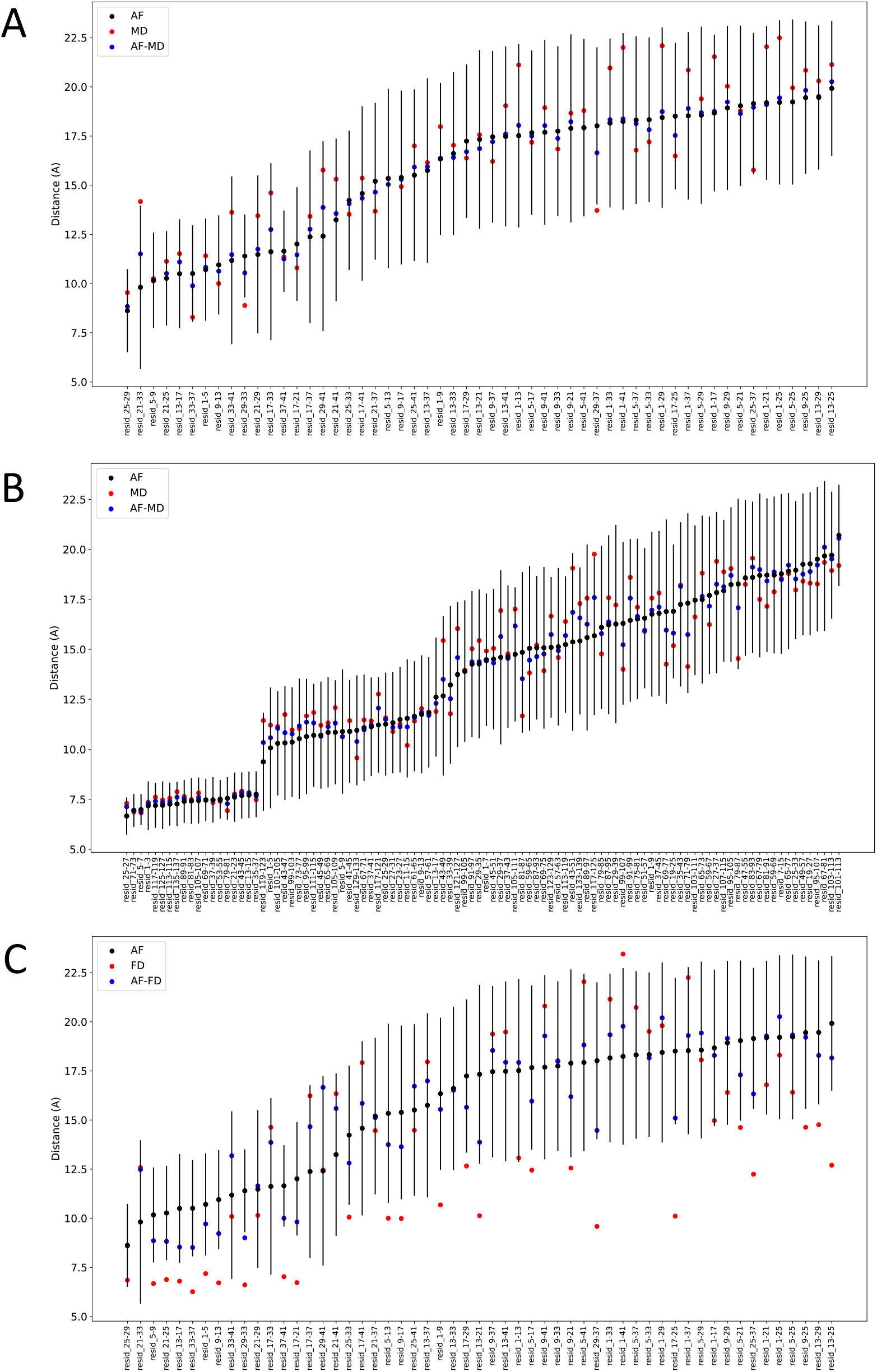
Comparison of inter-residue distances in the MD and AF-MD ensembles. (A) MD ensemble of Aβ. (B) MD ensemble of α-synuclein. (A) FD ensemble of Aβ. The inter-residue distances in the prior ensembles are shown as red points, the distances in the reweighted ensembles as blue points, and the AlphaFold distances are black points with an associated error bar (black line).

**Figure S2.**
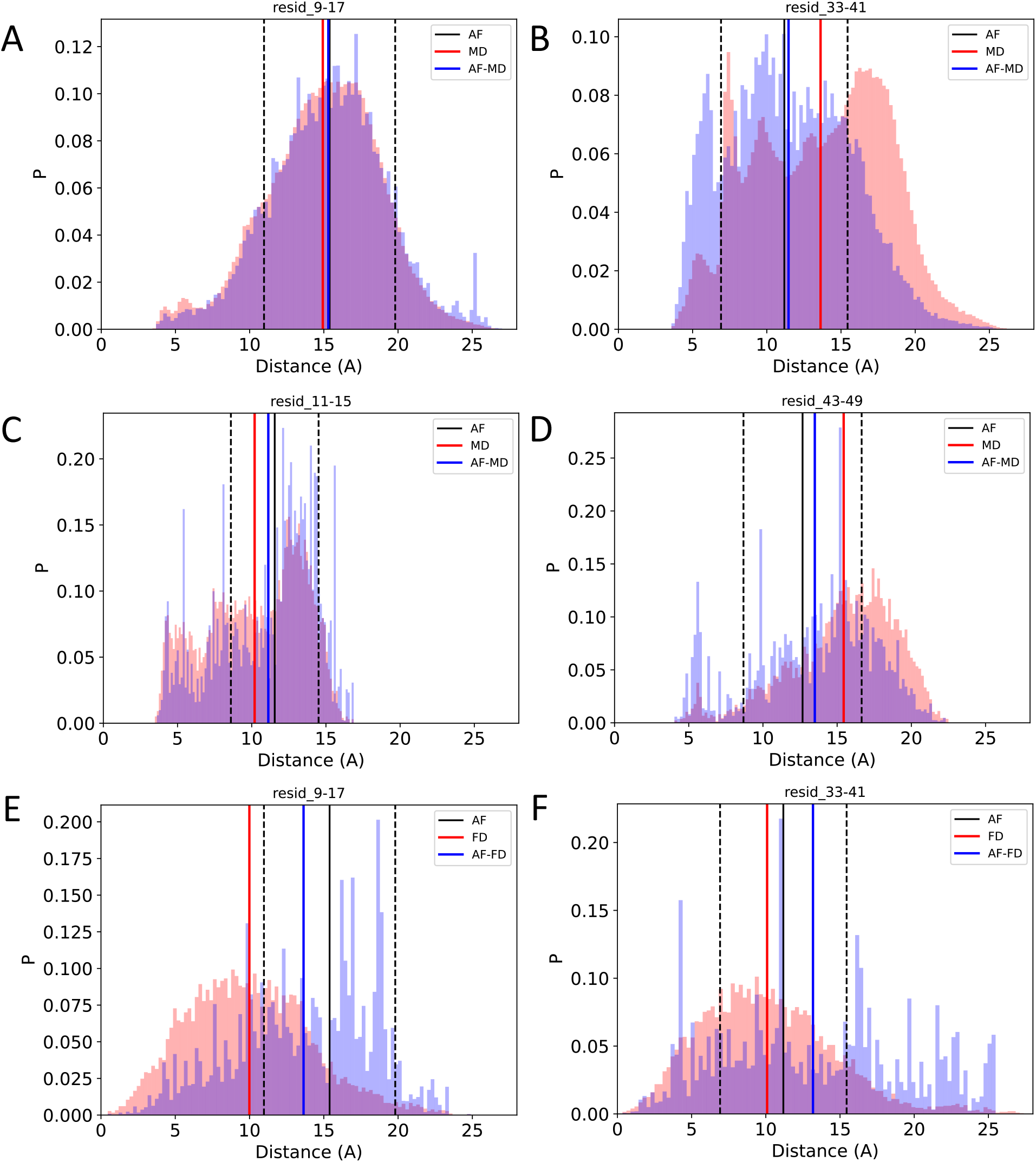
Comparison of selected inter-residue distance distributions in the MD, FD, AF-MD, and AD-FD ensembles of Aβ and α-synuclein. **(A**,**B)** MD ensemble of Aβ. **(C**,**D)** MD ensemble of α-synuclein. **(E**,**F)** FD ensemble of Aβ. Distributions in the MD and FD ensembles and AF-MD and AF-FD ensembles are colored in pink and purple, respectively; the average values are shown as red and blue solid lines, while the average AF distances and errors as black solid and dotted lines.

**Figure S3.**
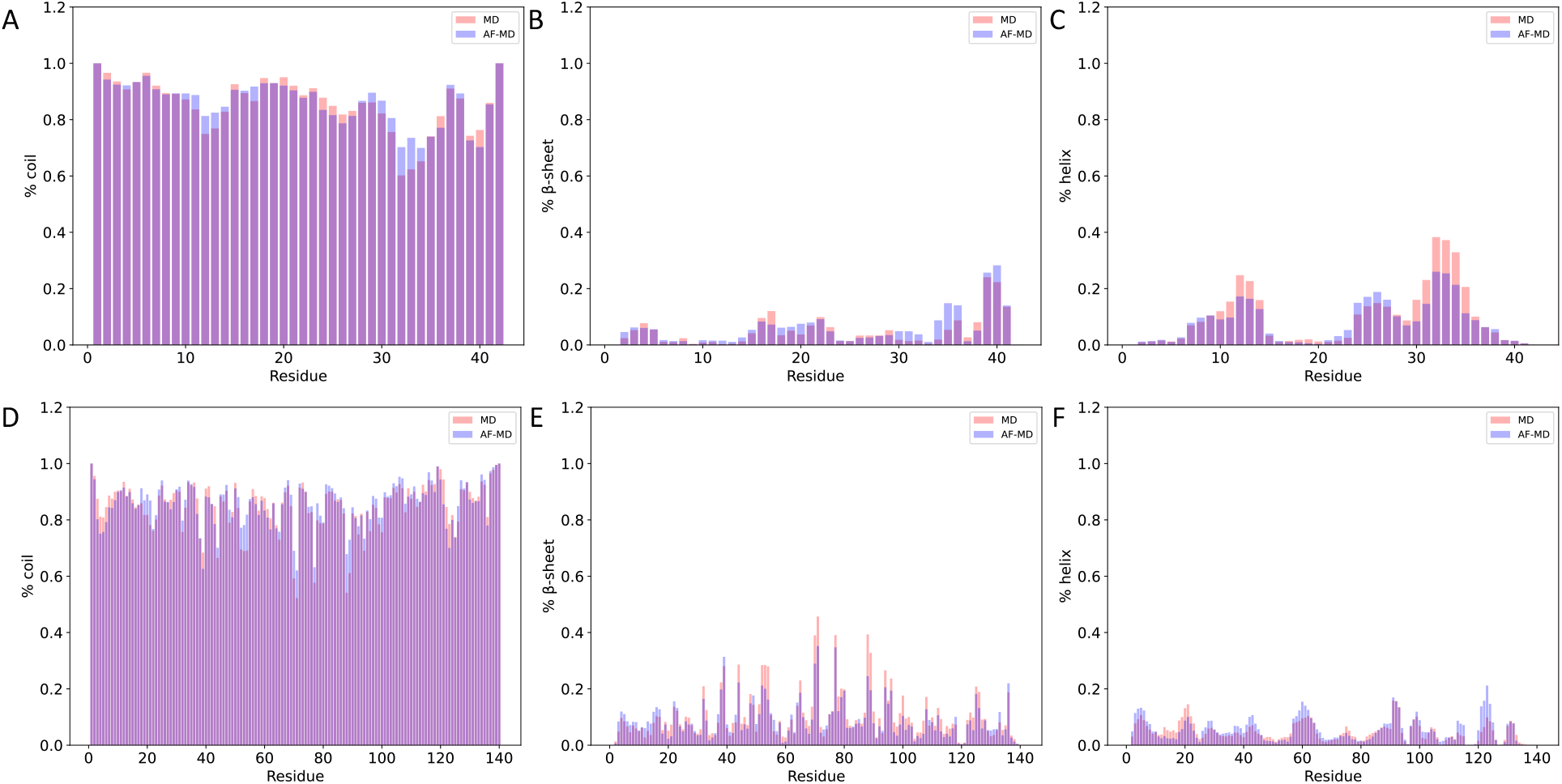
Comparison of secondary structure contents for the MD and AF-MD ensembles. **(A**,**D)** Random coil content of Aβ and α-synuclein. **(B**,**E)** β-sheet content of Aβ and α-synuclein. **(C**,**F)** α-helical content of Aβ and α-synuclein. MD and AF-MD ensemble predictions are shown in pink and purple, respectively.

